# Unequal interactions between alcohol and nicotine co-consumption: Suppression and enhancement of concurrent drug intake

**DOI:** 10.1101/601641

**Authors:** Margot C DeBaker, Janna K Moen, Jenna M Robinson, Kevin Wickman, Anna M Lee

## Abstract

**Rationale:** Alcohol and nicotine addiction are prevalent conditions that co-occur. Despite the prevalence of co-use, factors that influence the suppression and enhancement of concurrent alcohol and nicotine intake are largely unknown.

**Objectives:** Our goals were to assess how nicotine abstinence and availability influenced concurrent alcohol consumption, and to determine the impact of quinine adulteration of alcohol on aversion resistant alcohol consumption and concurrent nicotine consumption.

**Methods:** Male and female C57BL/6J mice voluntarily consumed unsweetened alcohol, nicotine and water in a chronic 3-bottle choice procedure. In Experiment 1, nicotine access was removed for 1 week and re-introduced the following week, while the alcohol and water bottles remained available at all times. In Experiment 2, quinine (100-1000 μM) was added to the 20% alcohol bottle, while the nicotine and water bottles remained unaltered.

**Results:** In Experiment 1, we found that alcohol consumption and preference were unaffected by the presence or absence of nicotine access in both male and female mice. In Experiment 2a, we found that quinine temporarily suppressed alcohol intake and enhanced concurrent nicotine, but not water, preference in both male and female mice. In Experiment 2b, chronic quinine suppression of alcohol intake increased nicotine consumption and preference in female mice without affecting water preference, whereas it increased water and nicotine preference in male mice.

**Conclusions:** Quinine suppression of alcohol consumption enhanced the preference for concurrent nicotine preference in male and female mice, suggesting that mice compensate for the quinine adulteration of alcohol by increasing their nicotine preference.

## Introduction

Alcohol and nicotine are the two most commonly abused addictive drugs and the majority of alcohol dependent individuals are also dependent on nicotine (Batel et al., 1995; Falk et al., 2006; Miller and Gold, 1998). Persons co-dependent on alcohol and nicotine have more severe drug dependence symptoms such as greater craving and withdrawal signs, higher drug consumption, increased difficulty maintaining abstinence, and have higher mortality compared with persons dependent on alcohol or nicotine alone (Heffner et al., 2011; Hurt et al., 1996; King et al., 2009; Leeman et al., 2008; Marks et al., 1997). Despite the high prevalence and increased health consequences of alcohol and nicotine co-dependence, there are currently no FDA-approved drugs for the treatment of alcohol and nicotine co-dependence. In human studies, it is difficult to dissect the effects of alcohol and nicotine from the genetic and environmental influences that also contribute to overall drug taking. Several reviews have highlighted key knowledge gaps that have limited the development of new drugs, such as the need to better understand the neurobiology of co-dependence and the need for identification of drug targets that mediate both alcohol and nicotine dependence (Tarren and Bartlett, 2017; Van Skike et al., 2016). To achieve this, the development of a greater variety of animal models that reflect alcohol and nicotine co-use is necessary, as this will enable the identification how alcohol and nicotine influence co-consumption while controlling for genetics and environment.

Many animal models of alcohol and nicotine co-dependence utilize investigator administered drugs (Blomqvist et al., 1996; Hendrickson et al., 2009; Lê et al., 2000, 2003; Smith et al., 2002), and although these models allow for control of dose and timing, they do not permit the animal to voluntarily consume both drugs. A limited number of studies in rats have examined voluntary alcohol and nicotine intake using several routes of administration, such as intravenous self-administration (IVSA) of nicotine with operant oral consumption of alcohol (Lê et al., 2010, 2014), IVSA nicotine with oral alcohol consumption in a 2-bottle choice procedure (Maggio et al., 2018), operant intra-cranial self-administration of nicotine with operant alcohol consumption (Deehan et al., 2015), or operant intra-cranial self-administration of a mixture of alcohol and nicotine (Truitt et al., 2015). These studies provide valuable information, yet the procedures involve significant training as well as technical and surgical requirements. In contrast, 2-bottle choice studies are frequently used in rats and mice to assess alcohol or nicotine consumption (Lee and Messing, 2011; Lee et al., 2014; Locklear et al., 2012; Meliska et al., 1995; Powers et al., 2013; Simms et al., 2008). In these studies, the animals are individually housed with two fluid bottles, one containing drug and one water, and the animals are free to consume from both bottles. These bottle choice studies are less technically challenging and are more high-throughput compared with operant administration studies.

We have previously developed a novel 3-bottle choice co-consumption model where mice voluntarily consume unsweetened alcohol, unsweetened nicotine, and water from 3 separate drinking bottles (O’Rourke et al., 2016; Touchette et al., 2018). Using this model, we investigated the effects of forced alcohol abstinence and intermittent drug access, and the impact of pre-clinical drug treatment on concurrent alcohol and nicotine consumption in male and female C57BL/6J mice (O’Rourke et al., 2016; Touchette et al., 2018). We reported that after 3 weeks of chronically co-consuming alcohol and nicotine, forced alcohol abstinence resulted in an increase in concurrent nicotine consumption and preference in male and female C57BL/6J mice, suggesting that the mice compensated for the absence of alcohol by increasing their consumption of nicotine (O’Rourke et al., 2016). It is unclear if the reciprocal relationship between alcohol and nicotine is true, and in the current study one of our goals was to determine whether forced nicotine abstinence enhanced concurrent alcohol consumption in this model.

A prominent feature of alcohol use disorders (AUDs) in humans is consumption of alcohol despite adverse legal, health, economic, and societal consequences. The continued consumption of alcohol despite the addition of the bitter tastant quinine has been frequently used as a model of compulsive alcohol intake or aversion-resistant alcohol consumption in both rats and mice (Hopf et al., 2010; Hopf and Lesscher, 2014; Lei et al., 2016; Sneddon et al., 2019; Spanagel et al., 1996). Our second goal was to determine whether the alcohol-abstinence induced elevation of nicotine consumption that we previously reported in O’Rourke et al, 2016 could be produced by suppressing alcohol consumption instead of removing alcohol access. Moreover, quinine suppression of alcohol consumption has not yet been evaluated in a voluntary alcohol and nicotine co-consumption model.

Here, we report an unequal interaction between alcohol and nicotine, where alcohol consumption is unaffected by the presence or absence of nicotine access. In contrast, the addition of quinine temporarily suppressed alcohol preference and enhanced concurrent nicotine preference in both sexes. Chronic suppression of alcohol consumption with quinine produced a long-term enhancement of concurrent nicotine, but not water, preference in female mice, and enhanced both nicotine and water preference in male mice.

## Materials and Methods

### Animals and Reagents

12 male and 12 female C57BL/6J mice from The Jackson Laboratory (Sacramento, CA) acclimated to our facility for at least six days before beginning behavioral experiments at 55 days old. All mice underwent both experimental procedures. Mice were group housed in standard cages under a 12-h light/dark cycle until the start of behavioral experiments, after which they were individually housed. Food and water were freely available at all times. All animal procedures were in accordance with the Institutional Animal Care and Use Committee at the University of Minnesota, and conformed to NIH guidelines.

Alcohol (ethanol) (Decon Labs, King of Prussia, PA), and nicotine tartrate salt (Acros Organics, Thermo Fisher Scientific, Chicago, IL) were mixed with tap water to the concentrations reported for each experiment. The concentrations of nicotine are reported as free base, and nicotine solutions were not filtered or pH adjusted. The alcohol, nicotine and water bottles were unsweetened at all times. Quinine hydrochloride (Spectrum Chemical Manufacturing Corp, Gardena, CA) was added to the 20% alcohol bottle at concentrations of 100, 200, 500 and 1000 μM in Experiment 2.

### Experiment 1: Effects of nicotine abstinence on concurrent alcohol consumption

Mice were singly housed in custom cages that accommodated three drinking bottles (Ancare, Bellmore, NY) containing water, nicotine or alcohol at different concentrations. The concentrations for the first week consisted of 3% alcohol (v/v) in one bottle, 5 μg/mL nicotine in the second bottle and water in the third bottle. The concentrations for the second week were 10% alcohol and 15 μg/mL nicotine, and were 20% alcohol and 30 μg/mL nicotine for the third week. During the fourth week, the nicotine bottle was removed and the 20% alcohol and water bottles remained available. During the fifth week, all three bottles were again presented at the 20% alcohol and 30 μg/mL nicotine concentrations. Food was freely available and the mice were not fluid restricted at any time. The bottles were weighed every other day and the solutions refreshed every 3-4 days. The positions of the bottles were alternated after each weighing to account for side preferences. Mice were weighed once a week. Fluid evaporation and potential dripping were accounted for by the presence of a set of alcohol, nicotine and water bottles on an empty control cage. The weight of fluid loss from these bottles was subtracted from all bottle weights throughout the study.

### Experiment 2: The impact of quinine adulteration on concurrent alcohol and nicotine consumption

Immediately after completion of Experiment 1, the mice proceeded to Experiment 2 without a break in alcohol or nicotine consumption. In Experiment 2a, quinine (100 μM) was added to the 20% alcohol bottle for 3 days (acute phase), while the water and 30 μg/mL nicotine bottles remained unaltered. In Experiment 2b, the quinine concentration in the 20% alcohol bottle was increased to 200, 500 and 1000 μM, with each concentration presented for 6 days (chronic phase). The bottles were weighed and rotated every day and the mice were weighed once a week. Fluid evaporation and bottle dripping were controlled for by the presence of a set of bottles on an empty control cage. The weight of fluid loss from these bottles was subtracted from all bottles throughout the study.

### Statistical Analysis

The average daily alcohol (g/kg) and nicotine consumption (mg/kg) for each drug concentration were calculated based on the weight of the fluid consumed from the bottles, the density of the solution (for alcohol only), and the weight of the individual mouse. The percent preference for alcohol, nicotine, and water bottles was calculated as the weight of the fluid consumed from the bottle, divided by the summed weight of fluid consumed from all three bottles, multiplied by 100. All analyses were calculated using Prism 8.0 (GraphPad, La Jolla, CA). Data were tested for normality and variance. Repeated comparisons with two independent variables were analyzed using two-way repeated measures ANOVA followed by Sidak’s or Tukey’s multiple comparisons tests, and repeated comparisons across one independent variable were analyzed with one-way repeated measures ANOVA followed by Dunnett’s multiple comparisons tests.

## Results

### Experiment 1: Effect of forced nicotine abstinence on concurrent alcohol consumption

A schematic outline of our experiments is shown in **Figure 1**. In Experiment 1, we evaluated the effect of forced nicotine abstinence on concurrent alcohol consumption and preference. Male and female C57BL/6J mice had continuous access to alcohol (v/v), nicotine (μg/mL) and water for 3 weeks in a 3-bottle choice procedure, with drug concentrations of 3% alcohol and 5 μg/mL nicotine presented during week 1, 10% alcohol and 15 μg/mL nicotine during week 2 and 20% alcohol and 30 μg/mL nicotine presented during week 3. During week 4, the nicotine bottle was removed and mice had access to the 20% alcohol and water bottles. The nicotine bottle was re-introduced in Week 5. There was a significant time by sex interaction for alcohol consumption, with female mice consuming more average alcohol (g/kg/day) compared with male mice for Weeks 3-5 (F_sexXtime_(4,88)=6.6061, *P*=0.0002; Sidak’s post-hoc test between sex: Week 3 *P*<0.0001, Week 4 *P*=0.0005, Week 5 *P*=0.004, **Figure 2A-B**, note scales on Y-axes). Female mice also consumed more nicotine (mg/kg/day) compared with male mice during Weeks 3 and 5 (F_sexXtime_(3,66)=3.070, *P*=0.03; Sidak’s post-hoc test between sex: Week 3 *P*=0.006, Week 5 *P*=0.004, **Figure 2A-B**). There was a significant time by sex interaction for alcohol preference (F_sexXtime_(4,88)=2.928) although post-hoc tests did not identify a significant sex difference in alcohol preference at any week (**Figure 2C-D**). We did not identify a sex difference in nicotine preference (F_sexXtime_(3,66)=1.408, *P*=0.25; F_sex_(1,22)=0.689, *P*=0.42; F_time_(3,66)=7.631, *P*<0.001).

**Fig. 1.**
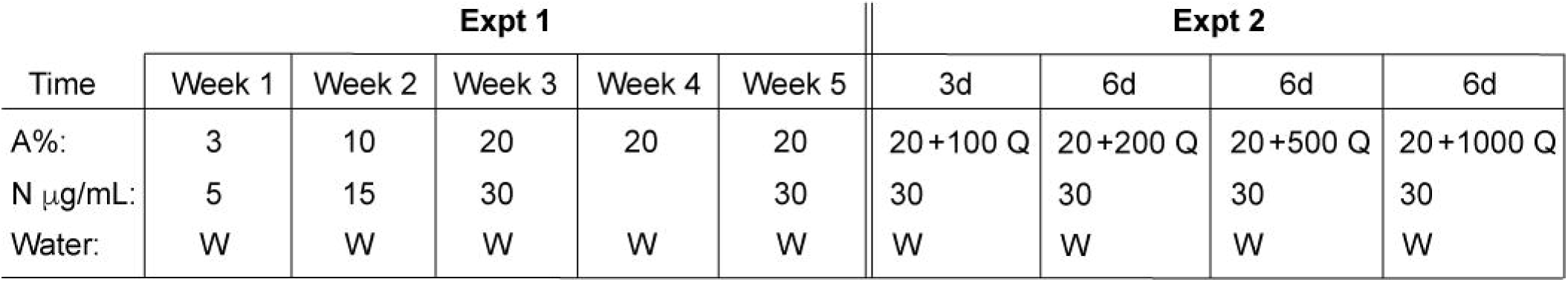
Schematic of experimental procedures. Unsweetened alcohol % (A, v/v), nicotine (N, μg/mL) and water were presented in a 3-bottle choice consumption procedure. Quinine (Q, μM) was added to the 20% alcohol bottle in Experiment 2

**Fig. 2.**
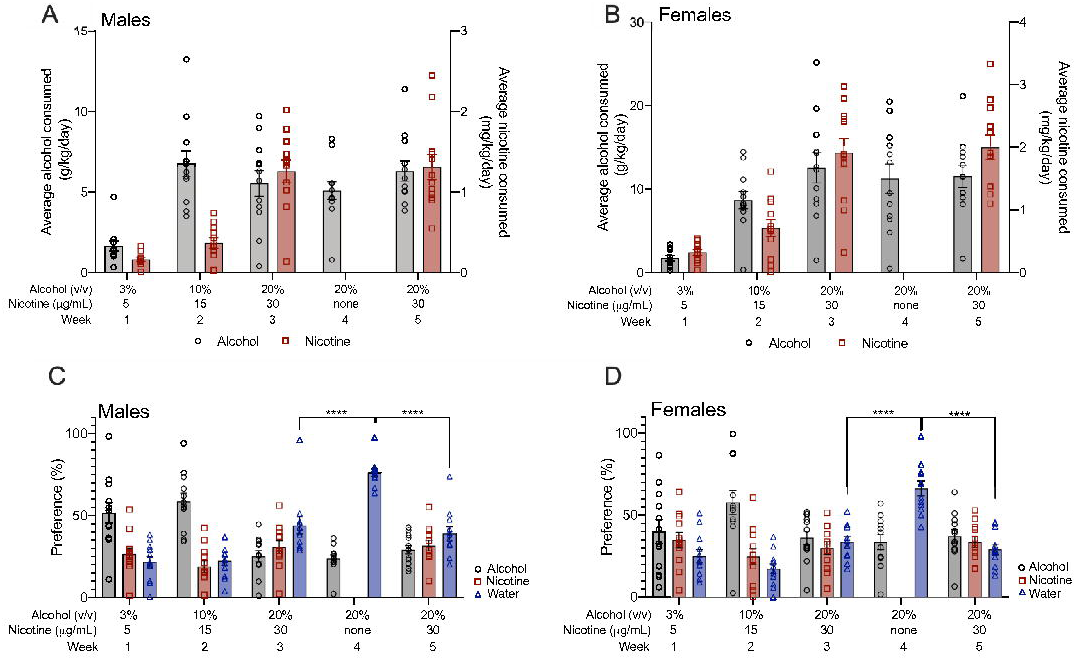
Nicotine forced abstinence increases concurrent water preference, and not alcohol preference, in male and female mice. Average alcohol (g/kg) and nicotine (mg/kg) consumption in (A) male and (B) female mice. Preference for the alcohol, nicotine and water bottles for (C) male and (D) female mice. Removal of the nicotine bottle during Week 4 increased the preference for the water bottle but not the alcohol bottle in (C) male and (D) female mice. Males: Tukey’s post-hoc test * * * * *P*<0.0001 between Weeks 3 and 4, and between Weeks 4 and 5. Females: Tukey’s post-hoc test * * * * *P*<0.0001 between Weeks 3 and 4, and between Weeks 4 and 5. *n*=12 per sex, mean ± SEM

During Week 4, we removed the nicotine bottle but maintained access to the alcohol and water bottles. We did not observe a change in the average alcohol consumption or alcohol preference across Weeks 3-5, indicating that removal of the nicotine bottle did not affect alcohol intake, though female mice still showed higher consumption and preference than male mice (alcohol consumption: F_sexXtime_(2,44)=0.549, *P*=0.58; F_time_(2,44)=0.622, *P*=0.54; F_sex_(1,22)=16.89, *P*<0.001; alcohol preference: F_sexXtime_(2,44)=0.253, *P*=0.78; F_time_(2,44)=2.320, *P*=0.11; F_sex_(1,22)=4.535, *P*=0.04). Interestingly, we found main effects of sex and time for water preference over Weeks 3-5 (F_sexXtime_(2,44)=0.002, *P*=0.99; F_time_(2,44)=107.2, *P*<0.0001; F_sex_(1,22)=5.181, *P*=0.03). Examining the water preference within sex showed that the preference for Week 4 was significantly higher in both males and females, suggesting that removal of the nicotine bottle elevated water but not alcohol preference (males: RM one-way ANOVA F(2,22)=75.73, *P*<0.0001; females: RM one-way ANOVA F(2,22)=41.58, *P*<0.0001; **Figure 2C-D**). We did not observe a significant change in nicotine consumption or preference after forced nicotine abstinence, suggesting that male and female mice readily return to their pre-forced abstinence nicotine intake (nicotine consumption: F_sexXtime_(1,22)=0.025, *P*=0.88; F_time_(1,22) =0.370, *P*=0.55; F_sex_(1,22)=8.251, *P*=0.009; nicotine preference: F_sexXtime_(1,22)=0.522, *P*=0.48; F_time_(1,22)=1.398, *P*=0.25; F_sex_(1,22)=0.01, *P*=0.91). Overall, Experiment 1 showed that forced nicotine abstinence increased the preference for water, but not alcohol, in both male and female mice.

### Experiment 2a: The impact of quinine adulteration on concurrent alcohol and nicotine consumption

We tested the effect of quinine adulteration of alcohol on concurrent alcohol and nicotine intake in male and female C57BL/6J mice. Quinine (100 μM) was added to the 20% alcohol bottle for 3 days. We found a significant sex by time interaction for alcohol consumption (F_sexXtime_(3,66)=4.142, *P*=0.009), with female mice consuming more alcohol compared with male mice for all days except for Day 1 (0 * * *P*=0.006, Day 2 * * *P*=0.004, Day 3 * *P*=0.04, **Figure 3A-B**, note scales on Y-axes). There was a main effect of time with no main effect of sex, or sex by time interaction for alcohol preference (F_sexXtime_(3,66)=2.415, *P*=0.07; F_time_(3,66)=35.99, *P*<0.0001; F_sex_(1,22)=0.87, *P*=0.36, **Figure 3C-D**). For nicotine consumption, there were main effects of sex and time without a significant interaction, with females consuming more nicotine compared with males (F_sexXtime_(3,66)=0.2655, *P*=0.85; F_time_(3,66)=3.367, *P*=0.02; F_sex_(1,22)=8.908, *P*=0.007). Similar to alcohol preference, there was a main effect of time with no effect of sex or significant interaction for nicotine preference (F_sexXtime_(3,66)=0.088, *P*=0.97; F_time_(3,66)=17.03, *P*<0.0001; F_sex_(1,22)=0.2834, *P*=0.60).

**Fig. 3.**
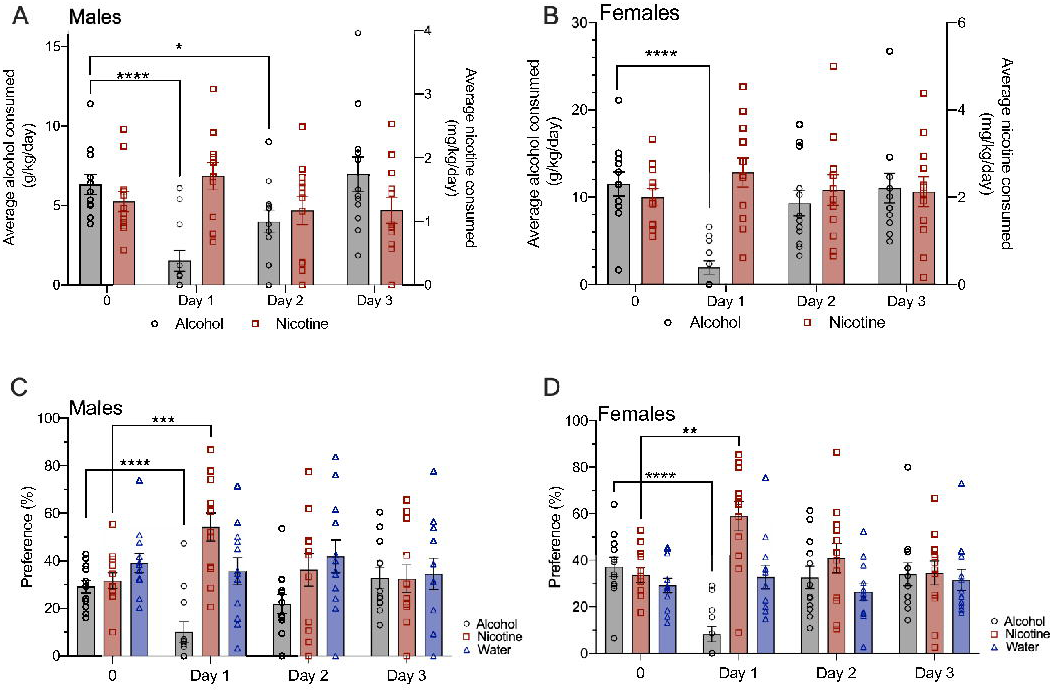
Temporary quinine-induced suppression of alcohol intake produces an increase in concurrent nicotine preference. Addition of 100 μM quinine to the 20% alcohol bottle occurred on Days 1-3. (A) Quinine suppressed alcohol consumption on Days 1 and 2 in male mice compared to the average alcohol consumption without quinine (0 time point). Dunnett’s multiple comparisons test * * * * *P*<0.0001 for Day 1, and * *P*=0.03 for Day 2 compared with 0. (B) Quinine suppressed alcohol consumption on Day 1 in female mice. Dunnett’s multiple comparisons test * * * * *P*<0.0001 for Day 1 compared with 0. (C) Quinine suppressed alcohol preference and increased nicotine preference on Day 1 in male mice. Dunnett’s multiple comparisons tests: alcohol preference * * * * *P*<0.0001 for Day 1 compared with 0, nicotine preference * * * *P*=0.0008 for Day 1 compared with 0. (D) Quinine suppressed alcohol preference and increased nicotine preference on Day 1 in female mice. Dunnett’s multiple comparisons tests: * * * * *P*<0.0001 for Day 1 compared with 0 for both alcohol and nicotine preference. *n*=12 per sex, mean ± SEM

We examined alcohol and nicotine consumption within sex and found that 100 μM quinine temporarily suppressed alcohol consumption in male mice on Days 1 and 2 (RM one-way ANOVA F(3,33)=15.83, *P*<0.0001, **Figure 3A**), and suppressed alcohol preference on Day 1 (RM one-way ANOVA F(3,33)=15.20, *P*<0.0001, **Figure 3C**). In female mice, 100 μM quinine adulteration of alcohol suppressed alcohol consumption on Day 1 (RM one-way ANOVA F(3,33)=22.16, *P*<0.0001, **Figure 3B**) and alcohol preference on Day 1 (RM one-way ANOVA F(3,33)=22.57, *P*<0.0001, **Figure 3D**). Quinine suppression of alcohol consumption and preference was temporary in both sexes, as both male and female mice overcame the 100 μM quinine suppression within 1-2 days.

The addition of 100 μM quinine to the 20% alcohol bottle produced a temporary increase in concurrent nicotine preference on Day 1 in male mice (RM one-way ANOVA F(3,33)=7.243, *P*=0.001, **Figure 3C**). Female mice showed a similar temporary increase in nicotine preference on Day 1 (RM one-way ANOVA F(3,33)=10.03, *P*<0.001, **Figure 3D**). There was a trend towards an increase in nicotine consumption on Day 1 in male mice (RM one-way ANOVA F(3,33)=2.649, *P*=0.07) but not in female mice (RM one-way ANOVA F(3,33)=1.358, *P*=0.27). There were no changes in water preference for either male or female mice (RM one-way ANOVAs, male: F(3,33)=0.704, *P*=0.56; female: F(3,33)=0.829, *P*=0.49).

### Experiment 2b: The impact of chronic quinine adulteration on concurrent alcohol and nicotine consumption

To determine the effect of chronic quinine adulteration of alcohol consumption on concurrent nicotine intake, the quinine concentration in the 20% alcohol bottle was increased to 200, 500 and 1000 μM, with each concentration presented for 6 days. We then evaluated the effect of quinine adulteration of alcohol across all quinine concentrations. There was a significant interaction between sex and concentration for alcohol consumption, with female mice consuming more alcohol compared with males for the 0, 100 and 200 μM quinine conditions (F_sexXconcentration_(4,88)=3.602, *P*=0.009; Sidak’s multiple comparisons test between sex: * * * * *P*<0.0001 for 0, * *P*<0.05 for 100 and 200 μM, **Figure 4A-B**, note scales on Y-axes). There was a main effect of concentration with no effect of sex, or a sex by concentration interaction for alcohol preference (F_sexXconcentration_(4,88)=0.958, *P*=0.43; F_concentration_(4,88)=52.54, *P*<0.0001; F_sex_(1,22)=1.318, *P*=0.26, **Figure 4C-D**). For nicotine consumption, there was a significant interaction between sex and concentration with female mice consuming more nicotine compared with male mice for the 200, 500 and 1000 μM quinine levels (F_sexXconcentration_(4,88)=5.156, *P*=0.0009; Sidak’s multiple comparisons test between sex: * * *P*<0.01 for 200 and 500 μM, * * * * *P*<0.0001 for 1000 μM). There was a significant interaction between sex and concentration for nicotine preference (F_sexXconcentration_(4,88)=2.835, *P*=0.03); however, multiple comparisons test did not reveal a significant difference in nicotine preference between sex at any concentration. We also found a main effect of sex and concentration with no interaction for water preference, with male mice having an overall greater preference for water compared with female mice (F_sexXconcentration_(4,88)=0.893, *P*=0.47; F_concentration_(4,88)=8.695, *P*<0.0001; F_sex_(1,22)=5.033, *P*=0.04, **Figure 4C-D**).

**Fig. 4.**
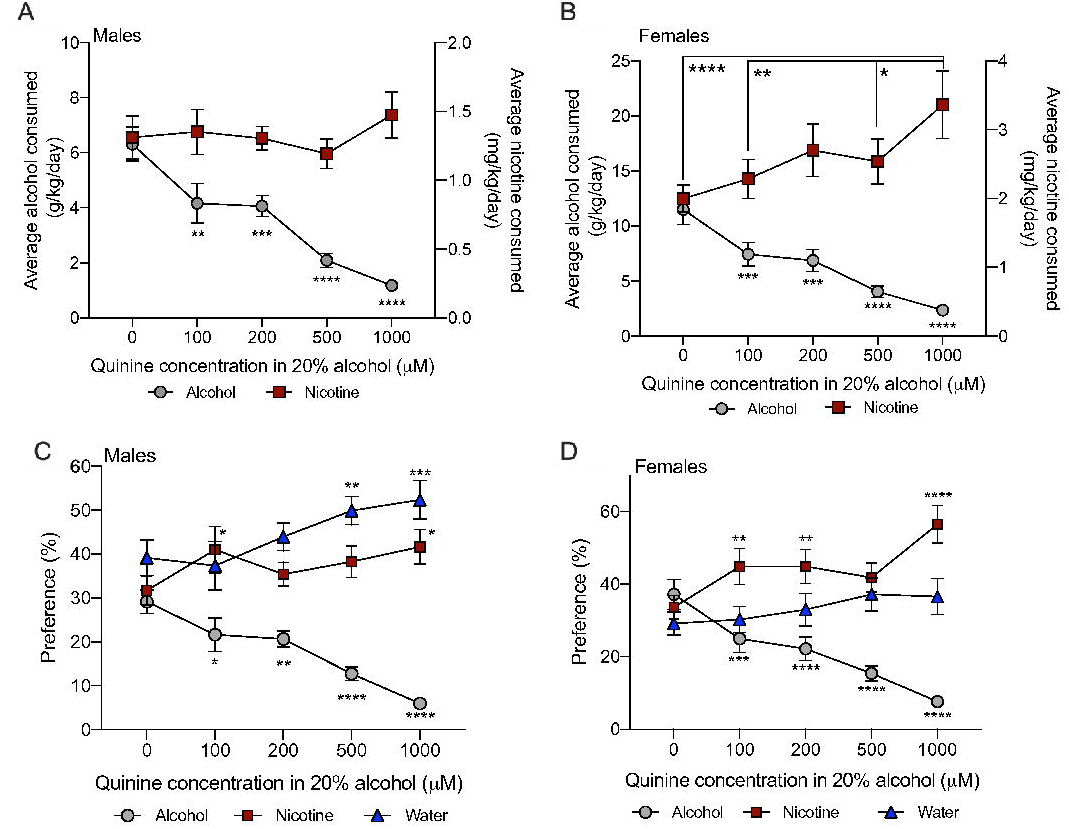
Chronic suppression of alcohol intake by quinine increases nicotine consumption and preference. (A) In male mice, increasing concentrations of quinine suppressed alcohol consumption without increasing nicotine consumption. (B) In female mice, increasing concentrations of quinine suppressed alcohol consumption and increased concurrent nicotine consumption. (C) Male mice showed reduced preference for the alcohol bottle at all concentrations of quinine. There was an increase in nicotine preference at the 100 and 1000 μM quinine concentrations, and an increase in water preference at the 500 and 1000 μM quinine concentrations compared with the same bottle at the 0 concentration. (D) Female mice showed a decrease in alcohol preference at all quinine concentrations. There was an increase in nicotine preference at 100, 200 and 1000 μM quinine concentrations, with no increase in water preference. All comparisons were Dunnett’s multiple comparisons tests at * *P*<0.05, * * *P*<0.01, * * * *P*<0.001, * * * * *P*<0.0001 compared with the 0 condition within drug bottle. *n*=12 per sex, mean ± SEM

Examining the effect of chronic quinine adulteration of alcohol consumption within sex, we found that the average daily alcohol consumption and preference was significantly suppressed at all quinine concentrations in male and female mice (RM one-way ANOVA, male alcohol consumption F(4,44)=25.48, *P*<0.0001; male alcohol preference F(4,44)=25.05, *P*<0.0001; female alcohol consumption F(4,44)=25.33, *P*<0.0001; female preference F(4,44)=27.98, *P*<0.0001). Chronic quinine adulteration of alcohol did not alter nicotine consumption over time in male mice (RM one-way ANOVA F(4,44)=1.310, *P*=0.28, **Figure 4A**), whereas female mice increased their nicotine consumption over time (RM one-way ANOVA F(4,44)=7.827, *P*<0.0001, **Figure 4B**). In male mice, nicotine preference was significantly increased when 100 and 1000 μM quinine was added to the 20% alcohol bottle, and water preference was significantly increased when 500 and 1000 μM quinine was added to the 20% alcohol bottle (F_drugXconcentration_(8,132)=14.34, *P*<0.0001, **Figure 4C**), showing that both nicotine and water preference increased during chronic quinine adulteration of alcohol. In contrast, female mice showed an increase in nicotine preference when 100, 200 and 1000 μM quinine was added to the 20% alcohol bottle, and did not show any significant increase in water preference at any quinine concentration (F_drugXconcentration_(8,132)=20.29, *P*<0.0001, **Figure 4D**), suggesting that female mice responded to chronic quinine adulteration of alcohol by consuming nicotine over water.

## Discussion

In this study, we used a 3-bottle choice procedure that allows for voluntary, chronic co-consumption of alcohol and nicotine to investigate the suppression and enhancement of concurrent drug intake in male and female C57BL/6J mice. We and others have published data showing that female mice consume significantly more drug compared with male mice (Hwa et al., 2011; Kamens et al., 2010, 2012; O’Rourke et al., 2016; Touchette et al., 2018), and here we also report significant sex differences in alcohol and nicotine consumption. In Experiment 1, we found that alcohol consumption and preference is unaffected by forced nicotine abstinence or the re-introduction of nicotine access in both male and female C57BL/6J mice. In Experiment 2a, we found that addition of 100 μM quinine to the 20% alcohol bottle temporarily suppressed alcohol consumption and preference, while increasing concurrent nicotine preference in male and female mice. In Experiment 2b, we found that chronic suppression of alcohol consumption with increasing concentrations of quinine increased both nicotine and water preference in male mice, and only increased nicotine preference in female mice. Our previous study showed that forced alcohol abstinence produced an enhancement in concurrent nicotine consumption and preference in male and female C57BL/6J mice (O’Rourke et al., 2016). Together with this study, our work showed that alcohol intake is unresponsive to the presence or absence of the nicotine bottle, but nicotine intake is influenced by the absence of alcohol consumption or by the adulteration of the alcohol bottle with quinine. In both studies, we used a 1-week forced abstinence period and it is possible that changes in concurrent alcohol consumption may require a longer abstinence duration.

One characteristic of AUDs is the continued consumption of alcohol despite adverse consequences. Quinine adulteration of alcohol consumption has been frequently used to model aversion-resistant alcohol intake (Hopf et al., 2010; Hopf and Lesscher, 2014; Lei et al., 2016; Sneddon et al., 2019; Spanagel et al., 1996). Quinine adulteration of alcohol consumption had not been previously examined in conjunction with concurrent nicotine consumption. In this study, we found that addition of 100 μM quinine to the alcohol bottle temporarily suppressed alcohol consumption and preference in both male and female mice. Interestingly, the suppression of alcohol intake was associated with increased preference for the nicotine bottle and not the water bottle. These data suggest that the mice compensate for the reduction in quinine-adulterated alcohol intake by selectively increasing preference for the unsweetened nicotine bottle, even though the water bottle is readily available. These results support our previous findings that nicotine consumption and preference is enhanced during forced alcohol abstinence (O’Rourke et al., 2016), and show that suppression of alcohol consumption, either by removing access to the alcohol bottle or reducing the palatability of alcohol, produces an enhancement of concurrent nicotine consumption.

The suppression of alcohol intake with 100 μM quinine was transient, as male and female mice overcame the aversion within 1-2 days. These data showed that mice chronically co-consuming alcohol and nicotine can overcome quinine adulteration of alcohol in a few days. Other recently published studies also show that C57BL/6J mice quickly develop quinine-resistant alcohol consumption. For example, Lesscher and colleagues (2010) show that male C57BL/6J mice are resistant to quinine adulteration of alcohol after 2 weeks of intermittent alcohol consumption (Lesscher et al., 2010), whereas Lei and colleagues (2016) show that a single session of unadulterated alcohol consumption is sufficient to produce aversion-resistant alcohol consumption (Lei et al., 2016). In contrast, prior studies in rats have required a minimum of 3 months of intermittent alcohol consumption before the development of aversion resistant alcohol consumption (Hopf et al., 2010; Spanagel et al., 1996).

We increased the concentration of quinine in the alcohol bottle to determine the effect of long-term suppression of alcohol consumption on concurrent nicotine intake. We found that alcohol consumption was suppressed in a concentration-dependent manner in both male and female mice. Male mice showed increased preference for the nicotine and water bottles as the quinine concentration increased, suggesting that male mice compensating for the long-term suppression of alcohol intake by increasing their nicotine and water preference. In contrast, female mice showed increased nicotine consumption and preference as the quinine concentration increased, and did not show any increase in water preference at any quinine concentration. Thus, our data suggest that female mice compensate for the long-term suppression of alcohol intake by increasing their nicotine intake. These data illustrate an important sex difference in compensatory drug consumption and highlight the need to continue investigating both sexes to identify important differences that can influence addiction-related behaviors and potential treatment strategies. Nearly all of the prior research on aversion-resistant drinking has focused on male animals. One recent study that investigates aversion-resistant drinking in male and female C57BL/6J mice shows no sex difference in the level of quinine suppressed alcohol consumption, even though female mice consume more alcohol than males (Sneddon et al., 2019). One limitation of our study is the lack of blood alcohol and nicotine levels. Since the mice are receiving access to the alcohol and nicotine bottles 24 hours a day, the consumption of alcohol and nicotine is spread out over time. Measuring blood alcohol and nicotine concentrations at an arbitrary time point is difficult due to the unsynchronized drug consumption and the fast metabolism of alcohol and nicotine in mice.

Alcohol and nicotine addiction are heritable disorders that share common genetic factors (Swan et al., 1996, 1997; True et al., 1999). The majority of tobacco smokers also use alcohol, yet smoking cessation trials frequently incorporate alcohol-related exclusion criteria and do not track co-use of alcohol in their subjects (Leeman et al., 2007), thus data on the treatment of co-users are lacking. Nearly one quarter of smokers report hazardous alcohol consumption patterns as defined by the NIAAA (Toll et al., 2012). These individuals have lower smoking cessation rates compared with moderate alcohol drinkers, highlighting that some co-users are less likely to successfully quit alcohol and nicotine use compared with other co-users (Toll et al., 2012). Varenicline (an α4β2 nicotinic acetylcholine receptor (nAChR) partial agonist) is approved for smoking cessation (Rollema et al., 2010), and naltrexone (an opioid receptor antagonist) and acamprosate (an NMDA receptor antagonist) are approved for alcohol use disorder (Franck and Jayaram-Lindström, 2013). Studies in alcohol-preferring female rats shows that varenicline reduces nicotine self-administration but has no effect on concurrent alcohol intake, and naltrexone reduces alcohol intake but has no effect on concurrent nicotine self-administration (Maggio et al., 2018; Waeiss et al., 2019), suggesting that monotherapy may be ineffective for combination alcohol and nicotine use disorder. Indeed, varenicline has shown mixed results in reducing alcohol consumption in human studies (de Bejczy et al., 2015; Litten et al., 2013; Mitchell et al., 2012; O’Malley et al., 2018; Plebani et al., 2013), and naltrexone and acamprosate fail to reduce cigarette smoking (Kahler et al., 2017; Fucito et al., 2012). Dual pharmacological treatment, such as combining naltrexone with nicotine replacement therapy, has been incorporated in trials to enhance the likelihood of successful alcohol and smoking abstinence (Kahler et al., 2017; Toll et al., 2010). Optimizing dual treatment strategies for different patterns of alcohol and nicotine co-use may provide further benefits. Moreover, pre-clinical research on combination alcohol and nicotine consumption will be necessary to understand the complex relationship between alcohol and nicotine co-use and identify novel drug targets that may be more helpful in treating human co-use.

In summary, our data highlight a complex interaction between alcohol and nicotine co-consumption. We found that the presence or absence of nicotine access does not affect alcohol consumption. However, adulterating alcohol with quinine produced a temporary suppression of alcohol consumption and preference, which was associated with an enhancement of concurrent nicotine preference. Chronic suppression of alcohol consumption with quinine produced compensatory nicotine preference in both male and female mice. Further behavioral and molecular dissection of these interactions will provide a better understanding of the neurobiology underlying alcohol and nicotine co-use.

